# Deep Learning-Based Analysis of Macaque Corneal Sub-Basal Nerve Fibers in Confocal Microscopy Images

**DOI:** 10.1101/758433

**Authors:** Jonathan D. Oakley, Daniel B. Russakoff, Megan E. McCarron, Rachel L. Weinberg, Jessica M. Izzi, Joseph L. Mankowski

## Abstract

**Purpose:** To describe and assess different deep learning-based methods for automated measurement of macaque corneal sub-basal nerves using in vivo confocal microscopy (IVCM).

**Methods:** The automated assessment of corneal nerve fiber length (CNFL) in IVCM images is of increasing clinical interest. These measurements are important biomarkers in a number of diseases including diabetes mellitus, human immunodeficiency virus, Parkinson’s disease and multiple sclerosis. Animal models of these and other diseases play an important role in understanding the disease processes as efforts toward developing new and effective therapeutics are made. And while automated methods exist for nerve fiber analysis in clinical data, differences in anatomy and image quality make the macaque data more challenging and has motivated the work reported here.

Toward this goal, nerves in macaque corneal IVCM images were manually labelled using an ImageJ plugin (NeuronJ). Different deep convolutional neural network (CNN) architectures were evaluated for accuracy relative to the ground truth manual tracings. The best performing model was used on separately acquired macaque ICVM images to additionally compare inter-reader variability.

**Conclusions:** Deep learning-based segmentation of sub-basal nerves in IVCM images shows excellent correlation to manual segmentations in macaque data. The technique is indistinguishable across readers and paves the way for more widespread adoption of objective automated analysis of sub-basal nerves in IVCM.

**Translational Relevance:** Quantitative measurements of corneal sub-basal nerves are important biomarkers for disease screening and management. This work reports on different approaches that, in using deep learning-based techniques, leverage state of the art analysis methods to demonstrate performance akin to human graders. In application, the approach is robust, rapid and objective, offering utility to a variety of clinical studies using IVCM.

## Introduction

In vivo confocal microscopy of the cornea (IVCM) allows for non-invasive acquisition of two-dimensional images, enabling detailed corneal sensory nerve fiber assessment in both clinical and research settings. The cornea is the most innervated tissue in the body, rendering the clinical applications of this non-invasive imaging procedure widespread. These applications include sensory neuropathy, where quantitative measurements of corneal sub-basal nerves are important biomarkers for disease screening and management. Measures of corneal sub-basal plexus nerve fiber count, density and length have been reported as having clinical utility in diabetes^1,2^, human immunodeficiency virus^3^, Parkinson’s disease^4^, multiple sclerosis^5^, and a number of other systemic illnesses. These metrics, however, are time consuming, require expertise, and are subjective when done manually. Automation of these measures is therefore necessary and will facilitate standardized analyses across centers as researchers investigate new end points in wide ranging clinical applications. As noted by Dabbah^6^, this lack of standardized assessment of corneal sub-basal nerve fiber density is a major limitation to wider adoption in clinical settings. Furthermore, the lack of a commonly accepted robust automated analysis method that provides centralized processing limits large-scale multicenter trials.

Several different approaches have been used to automate the task of nerve fiber tracing in IVCM. The challenging image conditions of noise, intensity hetereogeneity and low contrast features are compounded by the presence of dendritic, endothelial and inflammatory cells that have similar features to the nerves being delineated. Parissi7 adopted a graph-traversal method that traces between seed-points, which is an excellent way to describe the path of a nerve as the method effects constraints on feasible deviations of the nerve’s path and also bridges regions where the nerve’s intensity diminishes given the confocal nature of the modality. Fundamental to the success of such an approach is the choice of start and end points of the graph as these must belong to the same nerve. The original method for seed point detection is described by Scarpa^8^, where the image is covered in a grid of evenly spaced line-rows and columns. Nerves are detected at the intersection of lines based on intensity, and a tracing approach is used to follow the nerve in a direction perpendicular to its highest gradient. A final classification uses fuzzy c-means. Dabbah^6^ put an emphasis on carefully constructed filter banks applied as a feature detector and enhancing the nerves in accordance to their localized and dominant direction. A final binary image of nerves is created using a global threshold and skeletonization of the result. This work has resulted in the freely available ACCMetrics tool, widely accepted as a standard in IVCM image analysis^6,9,10,1,11,12^.

More advanced machine learning techniques have been added to the processing pipeline for a more sophisticated final arbitration between nerve fiber and background. Guimarães^13^ added a pixel-by-pixel classifier to hysteresis thresholded images to create a binarized nerve map. The features that fed the classifier were intensity based and included edge magnitudes. Structure enhancement used log-Gabor filters, and the method facilitates fast processing of large datasets. Annunziata^14^ developed a curvilinear structure model employing a set of filter banks to perform feature detection in the image. The parameters of the filters were hand tuned on the data set reported in the analysis. Additionally, contextual information is added via learned filters and a final classification then takes both of these results to yield an estimate at each pixel of nerve and background. The approach is reliant on both manual tuning and supervised learning for the feature designs, parameters and finally the thresholding. The results using cross-validation are impressive.

It can be seen that, in general, methods have evolved around being based on carefully designed filter banks acting as feature extractors with a final classification step. The best example of this is the ACCMetrics tool, which, being developed using human subject clinical data, offers a solution to the analysis of human corneal nerves acquired within a clinical environment. The methods, as implemented, should not, therefore, be expected to work “out of the box” on macaque data. As macaque models are used in a variety of diseases characterized by corneal sensory nerve fiber loss^15,16,17,18^, we have developed and characterized a novel approach for automated analysis of nerve fibers, leveraging more recent technologies in the world of computer vision and machine learning to process macaque IVCM images that are inherently of lower quality than human ICVM images.

Recent advances and superior levels of performance seen in the use of deep convolutional neural networks (CNNs) has resulted in their widespread adoption for a variety of image recognition tasks^19,20^. The deep-learning paradigm is to learn both the feature extraction (filters) and classifier using CNNs with supervised learning. The CNNs are capable of building rich, layered (deep) representations of the data which are then classified through additional layers of representation and learned associations. The following reports on using deep learning-based architectures for the automated tracing of corneal nerve fibers in IVCM images of macaque corneas.

## Materials and Methods

### Data

All data reported on in this study are derived from archived IVCM images acquired from anesthetized macaques using the Heidelberg HRTIII outfitted with the Rostock corneal module. In all cases, each image covered a field of view of 400μm by 400μm over 384 by 384 pixels. Using this information, the total lengths (mm) of the tracings can be converted to a measure of nerve length per image (mm/mm^2). This follows the convention of Dabbah^9^ in reporting the corneal nerve fiber length (CNFL), defined as the sum of the length of all nerves per image. For ground-truthing, sub-basal nerves were traced by experienced readers using the ImageJ plugin, NeuronJ^21^. Importantly, this is done at the pixel level, that is, the results of the labelling are images with each pixel labeled as belonging to either nerve or background.

### Methods

Based on the manually labelled data, a supervised deep-learning approach to semantic segmentation was used to associate the input images to this ground truth. This is done by presenting the network the image data and labels to create the pixel-wise associations. Categorical cross-entropy was used as the loss function that is minimized using backpropagation. The output of a trained network is then a nerve probability map where pixels are in the range 0, indicating no nerve data, to 1, just nerve data.

#### Pre-Processing and Post-Processing

Prior to presentation to the network a pre-processing step is used to account for differing background illumination across the image (Figure 1). This effect is all the more pronounced in macaque images given the increased curvature of the cornea in the macaques resulting in a faster roll-off of image intensities toward the periphery of the image than seen in human clinical data. Background illumination is first estimated by opening the image using grayscale morphology and a large structuring element, here a circle with a radius of 4μm (10 pixels). It then subtracts that result from the original image to reduce the changing background intensity. This is known as top hat filtering, where in this case, the brighter structures smaller than the structuring element are preserved.

**Figure 1.**
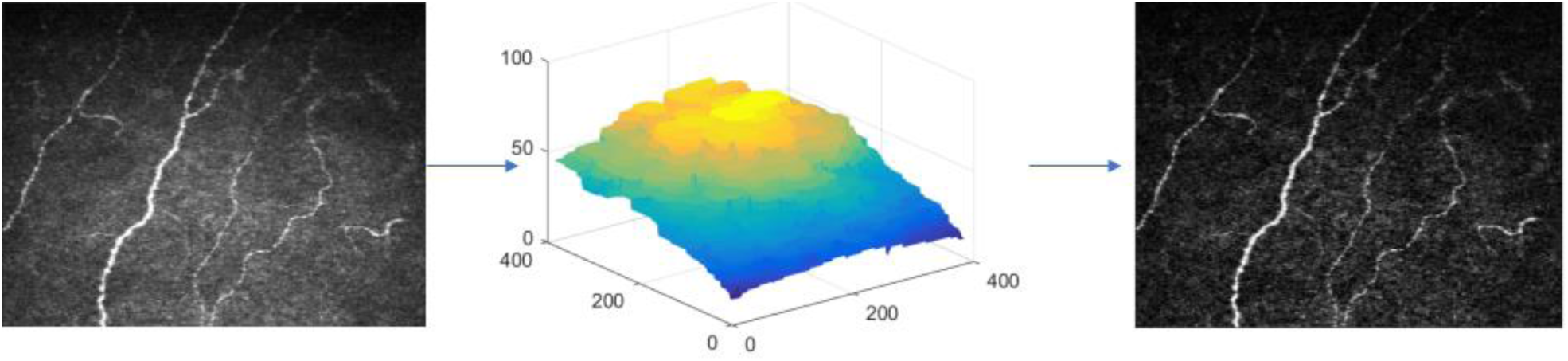
The pre-processing effects a simply flat-fielding of the image by estimating the background overall intensity using morphological opening (middle image) and subtracts that from the input image (left). The resulting image, on the right, has more even contrast across the field of view.

Post-processing is applied to the network’s output to threshold the probabilistic output, in the range 0 to 1, and return a final, binary result. Hysteresis thresholding with a lower value of 0.125 and an upper value of 0.275 are used for all cases in this study. The binary image is then skeletonized^22^ and components of less than 35 pixels are removed from consideration.

The combination of illumination correction on the front end and hysteresis thresholding on the back end work well in this application where the confocal nature of the imaging system mean that nerve fibers can come in and out of view (focus); that is, one must be locally sensitive and adaptive to the imaging conditions. The overall processing pipeline is illustrated in Figure 2, with the free parameters of the method being as given above; in the case of the flat-fielding, it is simply the size of the morphological structuring elements used to create the background image, and for post-processing it is the thresholds and the minimum individual component size. While the training stage can take hours, the final analysis takes, on average, two seconds per image.

**Figure 2.**
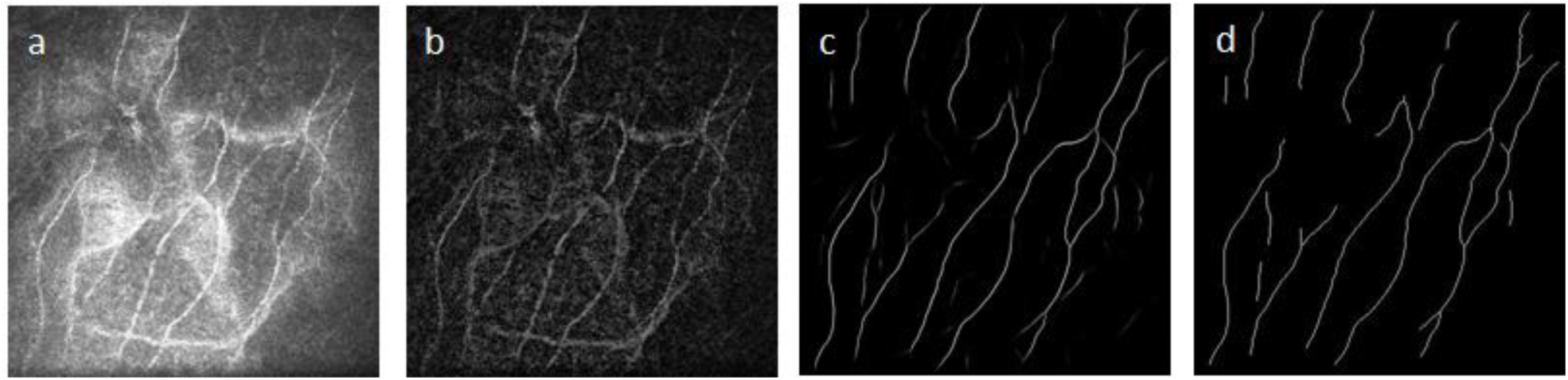
The input image (a) is pre-processed to compensate for variation in the background illumination (b). The segmentation, performed using deep learning, generates a probability image that assigns a score between 0 and 1 to each pixel (c) depicting a pixel-wise nerve classification. The final post-processing step is to binarize and skeletonize that result (d).

#### Neural Network Architectures

Three similar architectures for semantic segmentation were assessed, where candidate architectures were limited to networks that learned image to image mappings as a per-pixel classification. Common to the three architectures studied are encoding and decoding processing paths used to generate a final segmentation. Experiments involved altering the depth of the networks and also the optimization parameters, such as learning rate, decay and method as discussed in more detail below.

The first architecture is the autoencoder network^23^, as previously reported on^24^. We also looked at the U-net^25^, which is basically an auto-encoder network with additional connections across the encoding and decoding paths. The third is an extension of this method that adds skip connections within the encoding and decoding paths. This latter approach does this using residual branches that, in alleviating the vanishing gradient problem, facilitates deeper networks to be trained^26^. Performance is gauged using the coefficient of determination (R^2^) to assess strength of correlation between manual and automated results, as well as Bland-Altman analysis.

#### Deep Learning and Cross Validation

The different deep learning architectures were evaluated using 5-fold cross validation (CV), a standard approach for splitting train and test steps. The aforementioned pre- and post-processing steps were fixed for each as these admit to minimal parameterization as previously reported^27^. And while these parameters could also be learned using the CV paradigm, they were simple enough to tune manually once and left alone for all experiments.

In application of the CV, folds were chosen such that a single subject did not appear in both the training and test sets. For each fold, the training sets were additionally randomly split at each epoch into 90% training and 10% validation thereby allowing us to gauge how well the model’s learning was proceeding and when it should stop. This common practice is key in being able to decide on the optimizer used, batch sizes, as well as other hyper-parameters such as learning rate and learning decay as, in general, the loss value should decrease in a progressive way.

Results for all subjects and folds were pooled and compared to results using other architectures and hyper-parameters. For each of these experiments, a final correlation score between lengths reported by the manual tracings and those from the automated approach allowed us to rank the performance of the different implementations to derive the best model for each of the architectures used.

### Cross Validation Data and Results

This dataset comprised 58 IVCM images taken from 22 different macaques. To embellish the data, augmentation was used during the learning process. Summary correlations between the manual tracing and the automated approaches are given in Table 2 below, where we have included the result using ACCMetrics, the clinical tool applied to macaque data. Overall, the best performing architecture was the U-Net. Its configuration is given in Figure 4, where the optimizer used is the ADAM^28^ over 650 epochs; with the learning rate initialized at 10^−3^, dropping 5^-4^ every 10 epochs; a batch size of 8; and a drop-out rate of 0.2.

**Table 1.** Cross validation performance for the macaque data (N=58 from 22 subjects).

| Architecture | 5-Fold CV R<sup>2</sup> | Number of trainable parameters |
| --- | --- | --- |
| Autoencoder | 0.733 | 1,330,498 |
| U-Net | 0.859 | 487,730 |
| U-Net with Residual Branches | 0.731 | 516,578 |
| ACC Metrics | 0.718 | N/A |

**Table 2.** R^2^ correlations and intraclass correlation coefficients across all readers (ICCs are given in parenthesis). The higher the value, the more the agreement, overall, between the two readers. The ICC score across all readers was 0.84.

| Reader | R1 | R2 | R3 |
| --- | --- | --- | --- |
| R2 | 0.69 (0.84) |  |  |
| R3 | 0.82 (0.87) | 0.75 (0.80) |  |
| U-Net | 0.85 (0.85) | 0.69 (0.75) | 0.85 (0.92) |

**Figure 3.**
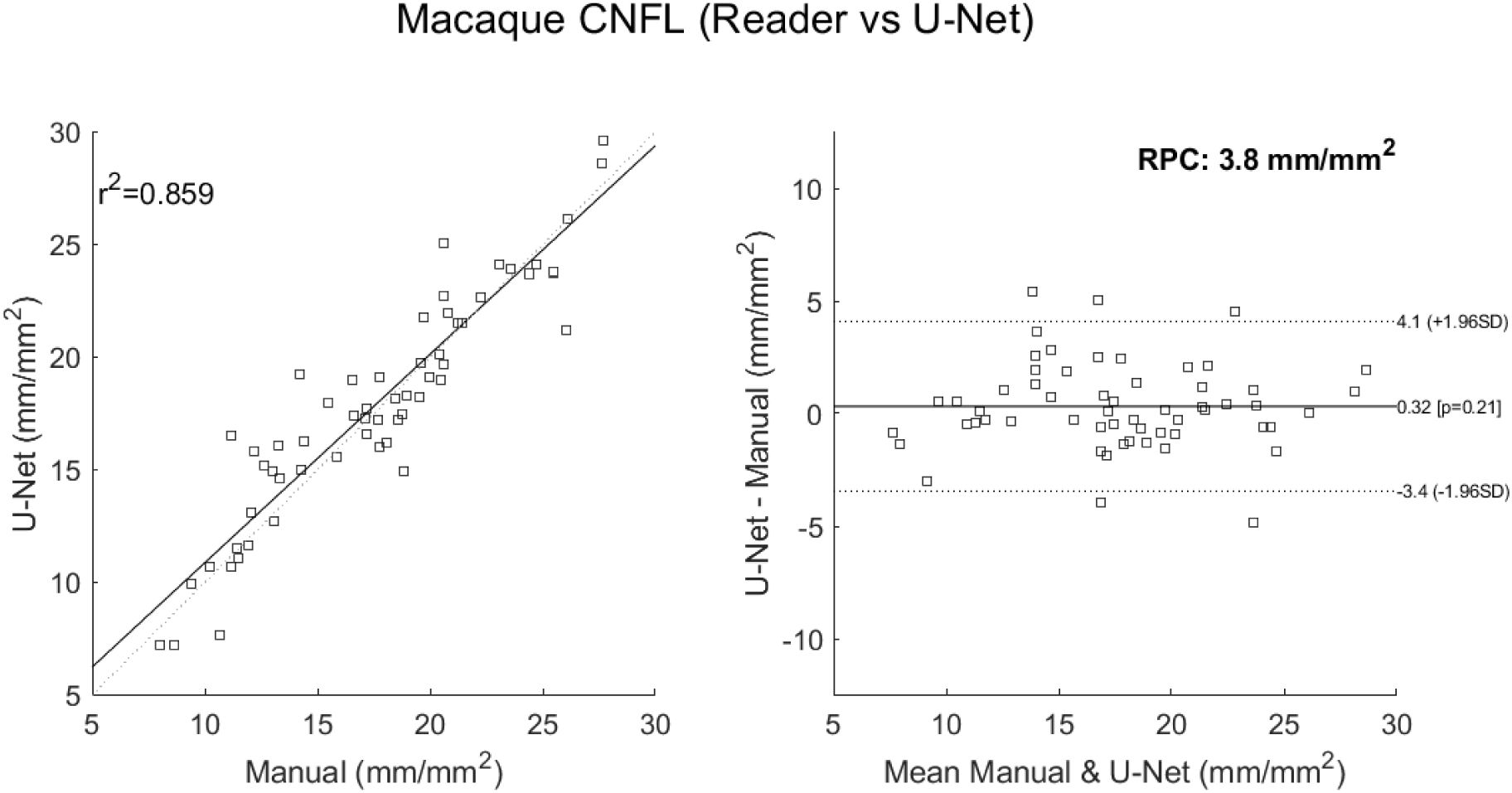
Correlations for the best performing U-Net result to the manual result for the macaque data using 5-fold CV (left). The limits of agreement show no systematic differences as the manual count increases or decreases. The reproducibility coefficient (RPC), 1.96*SD, is 3.8mm/mm^2^ (mean difference is 0.3mm/mm^2^).

**Figure 4.**
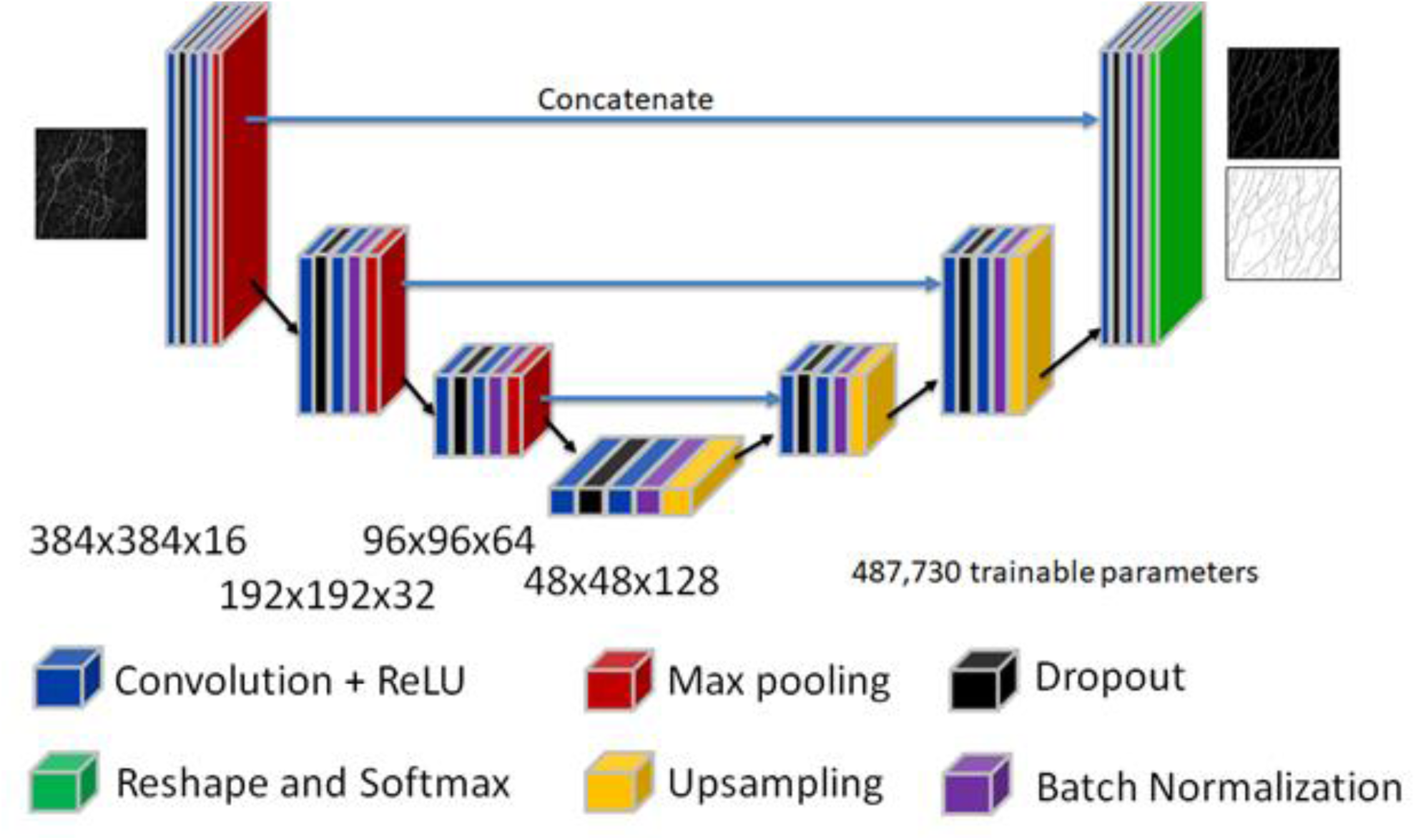
The best performing U-Net architecture used in these analyses. For the given input image, it outputs two images giving the probability score, at each pixel, of belonging to one of the two classes. The final softmax layer ensures these are normalized and can thus be interpreted as probabilities.

### Comparison Across Readers

The U-Net model that performed best in the macaque data CV analysis (Figure 3) was then trained on all folds; i.e., all 22 macaques in the data set. We then applied this model to a new population of 13 macaques that were being used in animal models of disease. In this instance, three readers were used to independently trace a total 46 images taken from these 13 subject macaques. This allows us to first see how well the best performing model can generalize to truly unseen data, and also to understand in such an applied environment how well it performs with respect to expert readers. Example manual and automated segmentation results are given in Figure 5.

In addition to the coefficient of correlation, intraclass correlation coefficients (ICCs) from two-way ANOVA analysis were derived to compare all readers and algorithm. The results are shown in Table 2 and all scores are good to excellent. Comparing average correlation scores for each reader relative to all other readers including the U-Net, yields R2s of: 0.79 (R1), 0.71 (R2), 0.81 (R3) and 0.80 (U-Net). By this metric, the automated method cannot be distinguished from the expert readers. This is confirmed by the ICC scores, where the score across all four observers is 0.84, which is exactly the same as the average U-Net ICC score across individual readers. It is worth noting here that the manual tracings that were used to train the network on the previously acquired data were from reader 1.

**Figure 5.**
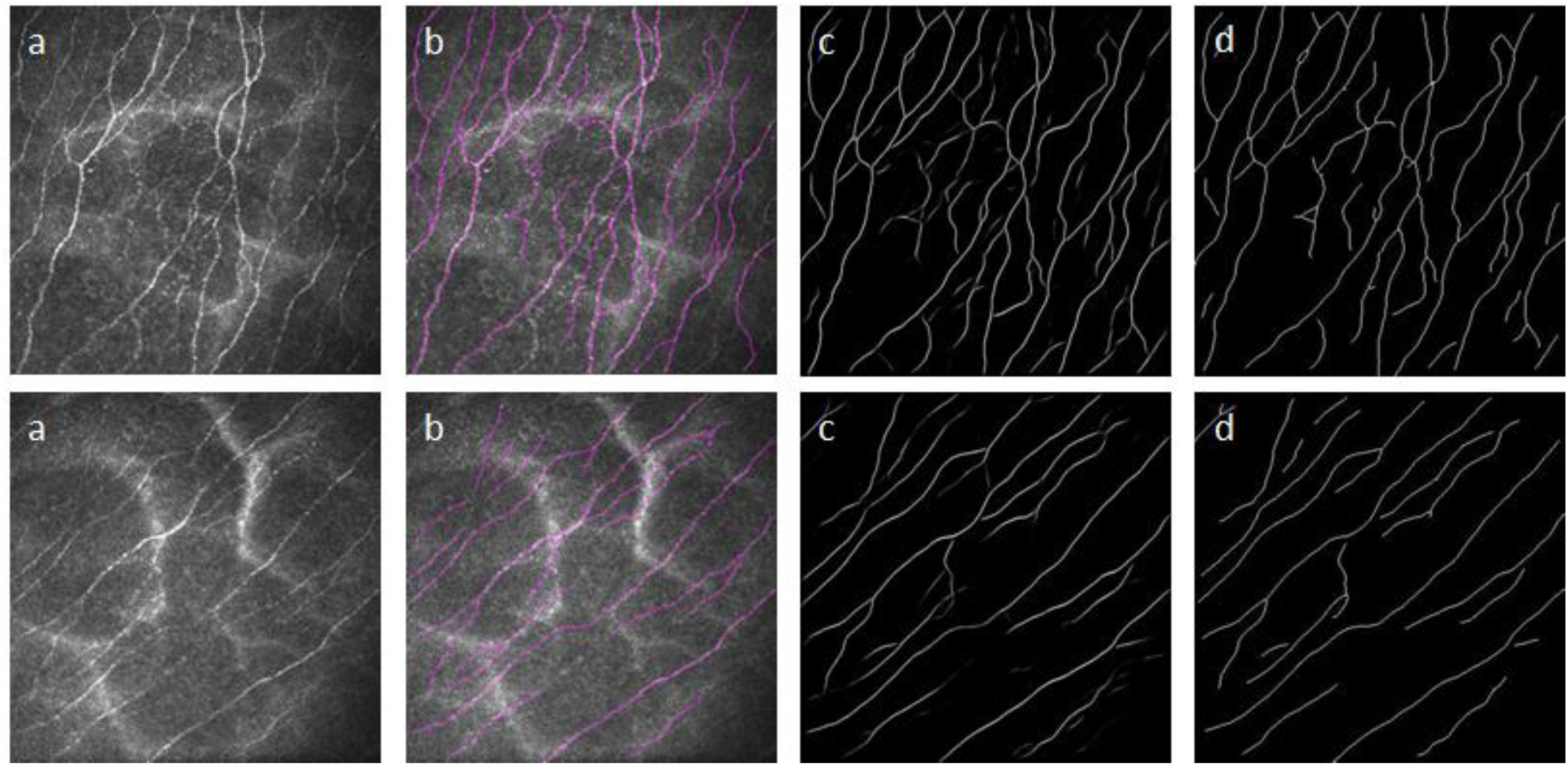
Example macaque data used during the inter-reader evaluation study. The images in the first column (a) are input images; the second column shows their manual tracings (b); the third column images are the probability images from the output of the neural network (c); and the final column gives those images thresholded and skeletonized (d).

### Comparison to the Average Reader

A final comparison looked at the correlation of two different methods to the average reader. These are:

- The U-Net model trained on 58 macaque images from 22 subjects applied to the new macaque data of 46 images (Figure 6). The LOA about the mean are given by the reproducibility coefficient (RPC) of 4mm/mm^2^.
- ACCMetrics applied to the new macaque data of 46 images (Figure 7). The RPC here increases to 4.5mm/mm^2^.

For both cases, the LOA are narrow, and the correlations are strong, with the U-Net being at 0.85 and ACCMetrics at 0.70.

**Figure 6.**
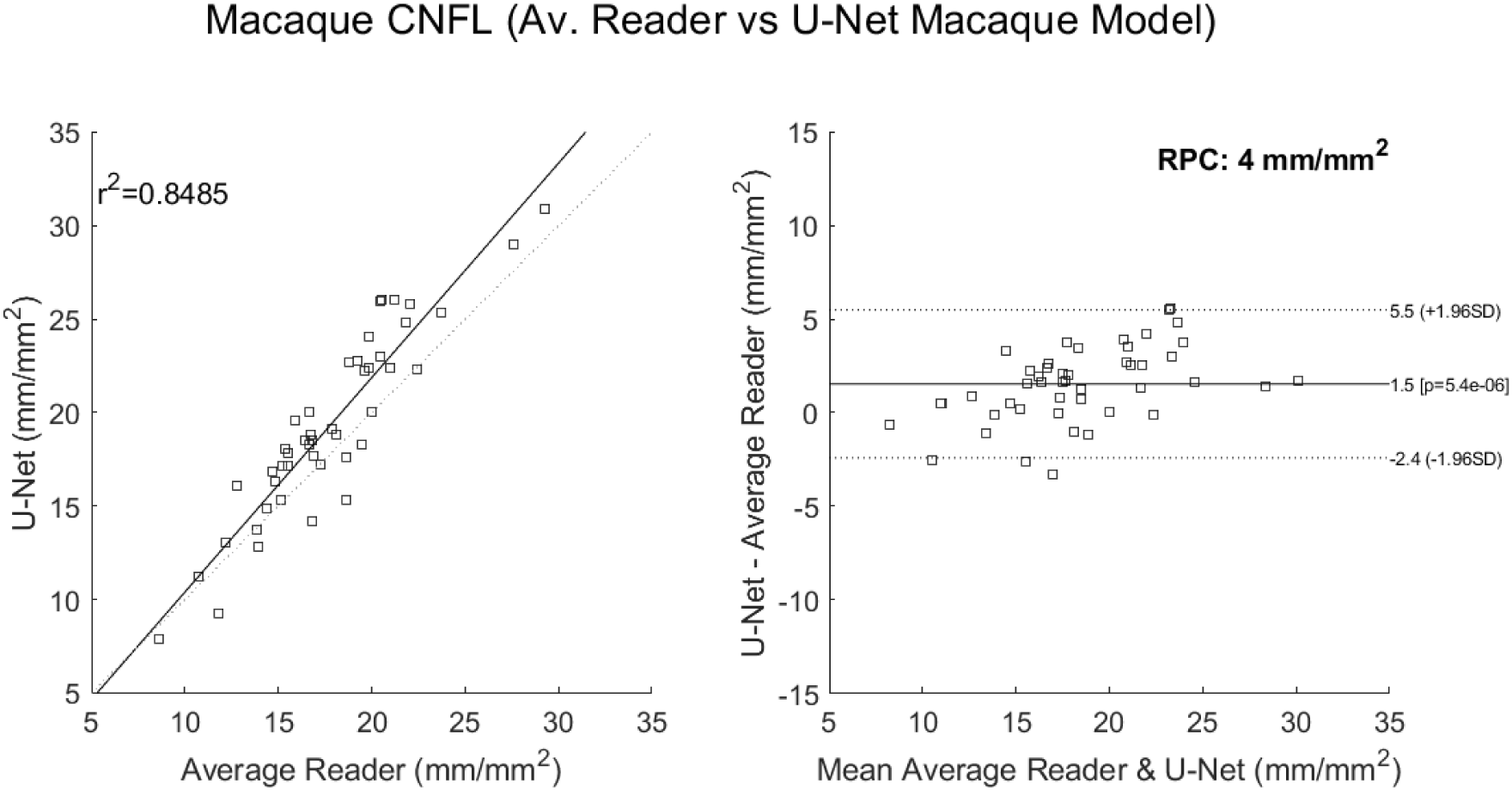
The above applies the best performing U-Net architecture using all of the macaque data (58 images) from the CV experiments. This is then applied to 46 images from 13 new subject macaques. This is our best performing method for the analysis of macaque data. The reproducibility coefficient (RPC), 1.96*SD, is 4mm/mm^2^ (mean difference is 1.5mm/mm^2^).

**Figure 7.**
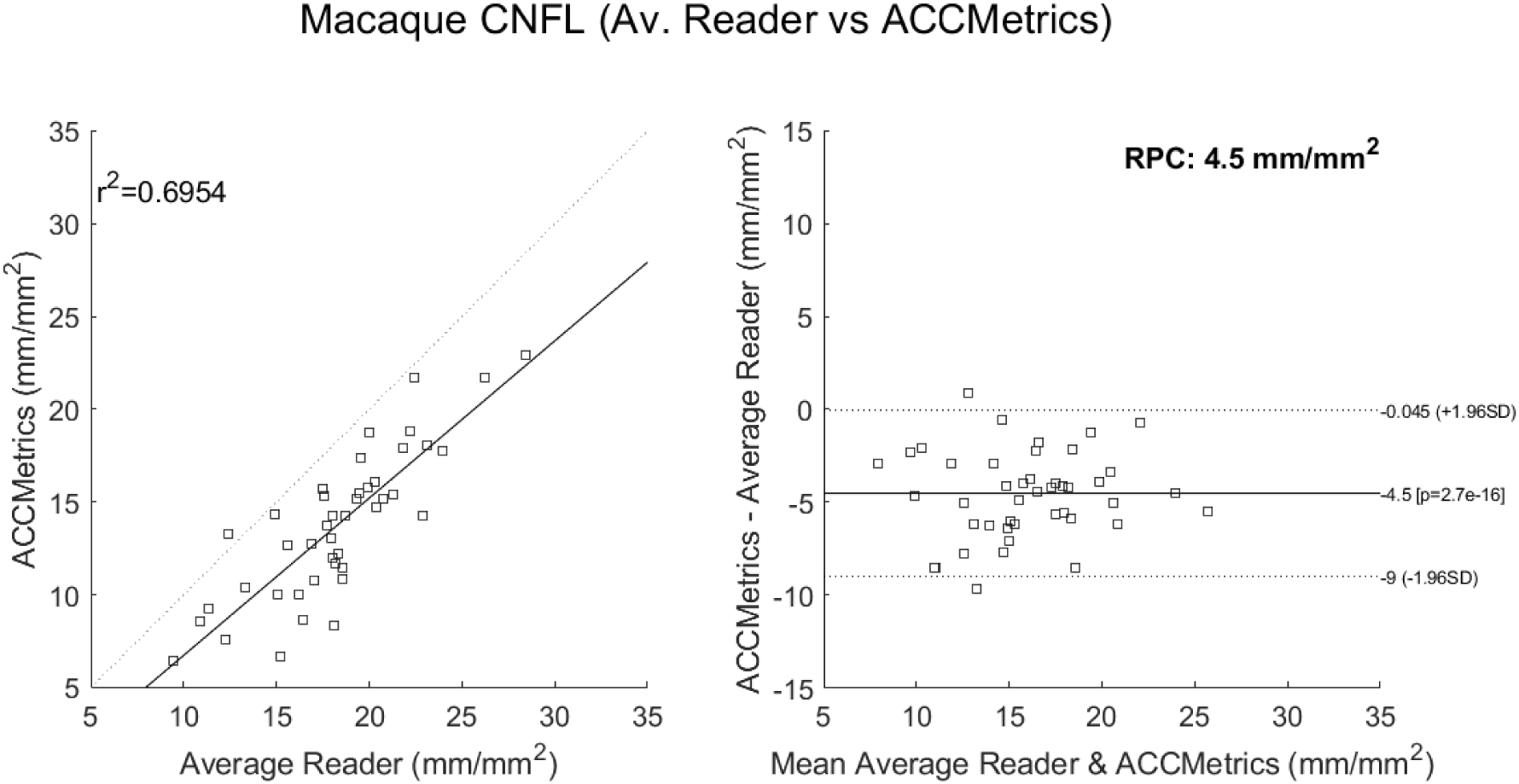
The above applies ACCMetrics to 43 of the 46 macaque images (three images did not process). Note that ACCMetrics was developed using clinical data, so the performance is expected to degrade using this data set. The reproducibility coefficient (RPC), 1.96*SD, is 4.5mm/mm^2^ (mean difference is −4.5mm/mm^2^).

## Discussion

There is wide-spread interest in using in vivo basal-nerve density assessment as a biomarker gauging corneal sensory nerve fiber loss. This is because of its relevance in a number of neuropathies and also systemic diseases. With such interest comes the need to automate the analysis, which, ahead of clinical adoption, also requires a validation of the accuracy of the approach. This study reports on a means of automating the analysis, here leveraging state of the art segmentation methods based on deep CNNs. It also presents an effort to validate the approach as applicable for use with macaque data. The method presented extends our original methods that were applied originally to ex vivo studies of immunstained corneal whole mounts^27^. Its extension to in vivo data and CCM imaging using deep learning was motivated by the research being done on animal models, but the work is clinically relevant if the reported performance extends to the use of clinically acquired data from human subjects. Furthermore, given a robust segmentation, derivative measures such as fractal density^29^ and tortuosity^14^ may also be investigated as biomarkers.

While comparison is made to ACCMetrics - a validated approach to clinical analysis of CCM images - it should here be emphasized that this software has not been developed for non-human analysis. This is an important difference as firstly the anatomy is different with a known increase in curvature of the cornea in the macaques and, in our review, macaque nerves may be generally thinner. We use ACCMetrics as a reference only, as it would be unfair to expect it to match the levels of performance reported in the literature when applied to macaque data. With that said, the performance is still good, has strong correlation to the manual readers, and fairly narrow limits of agreement in the Bland-Altman analysis. This serves, therefore, to speak to the overall robustness of the implementation. It also forewarns that, to apply our technique to clinical data, we will have to re-train the algorithm using just clinical data, as might be expected.

To conclude, the method we report on shows significant utility, here in the case of challenging macaque data. In conjunction with simple pre- and post-processing, excellent correlation with manual readings was achieved. In a comparison across readers, we have shown that the algorithm is indistinguishable from manual tracings. Lastly it should be noted that, while IVCM is currently the modality of choice, translation to other modalities should only require retraining of the neural network. There is, for example, increasing interest in the use of optical coherence tomography (OCT), the imaging standard of care in ophthalmology, for corneal nerve imaging^30^. If such a scenario evolves, this would make clinical adoption all the more likely in the future given the proliferation of devices and the ease of acquisition.

## References

1. Petropoulos IN, Alam U, Fadavi H, Marshall A, Asghar O, Dabbah MA, et al. Rapid automated diagnosis of diabetic peripheral neuropathy with in vivo corneal confocal microscopy. Investigative ophthalmology & visual science 2014; 55:2071–2078.

2. Misra SL, Craig JP, Patel DV, et al. In Vivo Confocal Microscopy of Corneal Nerves: An Ocular Biomarker for Peripheral and Cardiac Autonomic Neuropathy in Type 1 Diabetes Mellitus. Investigative ophthalmology & visual science 2015;56:5060–5065.

3. Kemp HI, Petropoulos IN, Rice ASC et al. Use of Corneal Confocal Microscopy to Evaluate Small Nerve Fibers in Patients With Human Immunodeficiency Virus. JAMA Ophthalmol. 2017 Jul 1;135(7):795–800

4. Misra SL, Kersten HM, Roxburgh RH et al. Corneal nerve microstructure in Parkinson’s disease. J Clin Neurosci. 2017 May;39:53–58.

5. Mikolajczak J, Zimmermann H, Kheirkhah A et al. Patients with multiple sclerosis demonstrate reduced subbasal corneal nerve fibre density. Mult Scler. 2017, Dec;23(14):1847–1853.

6. Dabbah MA, Graham J, Petropoulos I, Tavakoli M, Malik RA. Dual-model automatic detection of nerve-fibres in corneal confocal microscopy images, Med Image Comput Comput Assist Interv 13, 300-307, 2010.

7. Parissi M, Karanis G, Randjelovic S, et al. Standardized Baseline Human Corneal Subbasal Nerve Density for Clinical Investigations With Laser-Scanning in Vivo Confocal MicroscopyBaseline Corneal Subbasal Nerve Density. Investigative ophthalmology & visual science 2013;54:7091–7102.

8. Scarpa F, Grisan E, Ruggeri A. Automatic recognition of corneal nerve structures in images from confocal microscopy. Investigative ophthalmology & visual science 2008;49:4801–4807.

9. Dabbah MA, Graham J, Petropoulos I, Tavakoli M, Malik RA. Automatic Analysis of Diabetic Peripheral Neuropathy using Multi-scale Quantitative Morphology of Nerve Fibres in Corneal Confocal Microscopy Imaging, Journal of Medical Image Analysis 15(5), 738–747, 2011.

10. Petropoulos IN, Manzoor T, Morgan P, Fadavi H, Asghar O, Alam U, et al. Repeatability of in vivo corneal confocal microscopy to quantify corneal nerve morphology. Cornea 2013; 32:e83–89.

11. Chen X, Graham J, Dabbah MA, Petropoulos IN, Ponirakis G, Asghar O, et al. Small nerve fiber quantification in the diagnosis of diabetic sensorimotor polyneuropathy: comparing corneal confocal microscopy with intraepidermal nerve fiber density. Diabetes Care 2015; 38:1138–1144.

12. Chen X, Graham J, Dabbah MA, Petropoulos IN, Tavakoli M, Malik RA. An Automatic Tool for Quantification of Nerve Fibres in Corneal Confocal Microscopy Images. Chen, X. et al. (2016) IEEE Transactions on Biomedical Engineering, 2017 Apr;64(4):786–794.

13. Guimarães P, Wigdahl J, Ruggeri A. A Fast and Efficient Technique for the Automatic Tracing of Corneal Nerves in Confocal Microscopy. Translational Vision Science & Technology 2016;5.

14. Annunziata R, Kheirkhah A, Aggarwal S, Hamrah P, Trucco E. A fully automated tortuosity quantification system with application to corneal nerve fibres in confocal microscopy images. Medical image analysis 2016;32:216–232.

15. Laast VA, Pardo CA, Tarwater PM et al. Pathogenesis of simian immunodeficiency virus-induced alterations in macaque trigeminal ganglia. J Neuropathol Exp Neurol. 2007 Jan;66(1):26–34.

16. Paré M, Albrecht PJ, Noto CJ et al. Differential hypertrophy and atrophy among all types of cutaneous innervation in the glabrous skin of the monkey hand during aging and naturally occurring type 2 diabetes. J Comp Neurol. 2007 Apr 1;501(4):543–67.

17. Laast VA, Shim B, Johanek LM et al. Macrophage-mediated dorsal root ganglion damage precedes altered nerve conduction in SIV-infected macaques. Am J Pathol. 2011 Nov;179(5):2337–45.

18. Dorsey JL, Mangus LM, Hauer P et al. Persistent Peripheral Nervous System Damage in Simian Immunodeficiency Virus-Infected Macaques Receiving Antiretroviral Therapy. J Neuropathol Exp Neurol. 2015 Nov;74(11):1053–60.

19. Krizhevsky, A., Sutskever, I., and Hinton, G. E. (2012). Imagenet classification with deep convolutional neural networks. In Pereira, F., Burges, C., Bottou, L., and Weinberger, K., editors, Advances in Neural Information Processing Systems 25, pages 1097–1105, 2012.

20. Russakovsky, O., Deng, J., Su, H., Krause, J., Satheesh, S., Ma, S., Huang, Z., Karpathy, A., Khosla, A., Bernstein, M., Berg, A., and Fei-Fei, L. (2015). Imagenet large scale visual recognition challenge. International Journal of Computer Vision, 115(3):211–252.

21. Meijering E, Jacob M, Sarria JC, Steiner P, Hirling H, Unser M. Design and validation of a tool for neurite tracing and analysis in fluorescence microscopy images. Cytometry Part A 2004;58:167–176.

22. Zhang TY and Suen CY. A fast parallel algorithm for thinning digital patterns. Communications of the ACM, March 1984, Volume 27, Number 3.

23. Badrinarayanan 2016 – Badrinarayanan V, Kendall A, Cipolla R. SegNet: A Deep Convolutional Encoder-Decoder Architecture for Image Segmentation, 1511.00561.

24. Oakley JD, Russakoff DB, Weinberg R et al. Automated Analysis of In Vivo Confocal Microscopy Corneal Images Using Deep Learning. Investigative Ophthalmology & Visual Science July 2018, Vol.59, 1799.

25. Ronneberger 2015 - Ronneberger O, Fischer P and Brox T. Medical Image Computing and Computer-Assisted Intervention (MICCAI), Springer, LNCS, Vol.9351: 234-241, 2015.

26. He K, Zhang X, Ren S and Sun J. Deep Residual Learning for Image Recognition, 2016 IEEE Conference on Computer Vision and Pattern Recognition (CVPR).

27. Dorsey JL, Mangus LM, Oakley JD et al. Loss of Corneal Sensory Nerve Fibers in SIV-Infected Macaques: An Alternate Approach to Investigate HIV-Induced PNS Damage. Am J Pathol. 2014 Jun; 184(6):1652–1659.

28. Kingma DP and Adam BJ. A Method for Stochastic Optimization. (2014) cite 1412.6980.

29. Chen X, Grahnam J, Petropoulus IN et al. Corneal Nerve Fractal Dimension: A Novel Corneal Nerve Metric for the Diagnosis of Diabetic Sensorimotor Polyneuropathy. Invest Ophthalmol Vis Sci. 2018 Feb 1;59(2):1113–1118.

30. Hosseinaee Z, Han L, Kralj O et al. Fully automated segmentation algorithm for corneal nerves analysis from in-vivo UHR-OCT images. Investigative Ophthalmology & Visual Science July 2019, Vol.60, 167.

